# Layer 5 circuits in V1 differentially control visuomotor behavior

**DOI:** 10.1101/540807

**Authors:** Lan Tang, Michael J. Higley

## Abstract

Sensory areas of the mammalian neocortex are thought to act as distribution hubs, transmitting information about the external environment to various cortical and subcortical structures in order to generate adaptive behavior. However, the exact role of cortical circuits in sensory perception remains unclear. Within primary visual cortex (V1), various populations of pyramidal neurons (PNs) send axonal projections to distinct targets, suggesting multiple cellular networks that may be independently engaged during the generation of behavior. Here, we investigated whether PN subpopulations differentially support sensory-guided performance by training mice in a visual detection task based on eyeblink conditioning. Applying 2-photon calcium imaging and optogenetic manipulation of anatomically-defined PNs in behaving animals, we show that layer 5 corticopontine neurons strongly encode task information and are selectively necessary for performance. These results contrast with recent observations of operant behavior showing a limited role for the neocortex in sensory detection. Instead, our findings support a model in which target-specific cortical subnetworks form the basis for adaptive behavior by directing relevant information to downstream brain areas. Overall, this work highlights the potential for neurons to form physically interspersed but functionally segregated networks capable of parallel, independent control of perception and behavior.

## Introduction

The role of the mammalian neocortex in generating behavior in response to external sensory information has been investigated in mice across a range of modalities, including visual, auditory, and somatosensory inputs^1–7^. In some cases, inactivation of cortical circuits may lead to a disruption of task performance^2,4,5,7^. However, recent work has suggested that the detection of sensory cues linked to behavior may not require cortical function, even for optimal performance levels^3^.

The reasons for these discrepancies are not entirely clear, but likely stem from differences in the nature of the sensory stimulus, the required motor response, and the specific neuronal circuits engaged across different experimental paradigms. Thus, the necessity for cortical involvement in a particular task may be wholly dependent on the specific sensory and motor pathways, and their constituent microcircuits, engaged during that behavior. Consistent with this view of compartmentalized neural organization, there is a growing appreciation for the diversity of cell types in the neocortex, with categories determined by molecular and genetic markers, electrophysiological properties, morphology, and projection patterns^8–10^. Multiple *ex vivo* studies have provided evidence that this diversity extends to patterns of connectivity both within and across layers, suggesting independent, parallel networks may be common^11–15^. Moreover, investigations of somatosensory, auditory, and visual cortical areas have demonstrated that projection-specific subtypes of excitatory pyramidal neurons (PNs) can differentially encode information about sensory stimuli and motor action^4,16–19^ suggesting segregated streams of information may be directed to various cortical and subcortical targets. However, the functional independence of these physically interspersed local circuits during perception and sensorimotor behavior remains unclear.

The question of microcircuit independence is particularly relevant in layer 5, the principal output from the neocortex to subcortical structures. There, the two primary cell categories comprise neurons that target either the superior colliculus and brainstem or other intratelencephalic areas including the ipsilateral cortex and striatum^10,20–24^. We previously showed that under anesthesia, brainstem-projecting PNs in mouse V1 show higher contrast sensitivity and broader tuning for orientation and spatial frequency in comparison with striatal-projecting cells, suggesting the possibility that these largely non-overlapping populations may form functionally independent microcircuits^17^. Nevertheless, it has remained untested whether such distinct networks might be differentially engaged in and required for visually-guided behavior.

To address this question, we developed a novel detection task based on visually-cued eyeblink conditioning^25^, requiring the animal to detect a small drifting grating stimulus in order to perform a conditioned blink and avoid a corneal air puff. We first show that this task requires V1 function for both acquisition and performance of contrast-dependent behavior. Then, we apply an intersectional viral labeling strategy to express GCaMP6s in targeted subpopulations of layer 5 PNs and use 2-photon imaging of neuronal activity to show that corticopontine (CPn) but not corticostriatal (CSt) cells robustly encode information about individual trial performance in their stimulus-evoked responses. Finally, we show that optogenetic suppression of CPn activity, but not CSt activity, disrupts task performance. Overall, these data provide strong evidence for a cortical contribution to sensory detection and demonstrate the functional independence of microcircuits in the primary output layer of V1.

## Results

To investigate the functional architecture of mouse primary visual cortex (V1) underlying visual perception, we developed a novel detection task based on visually-cued eyeblink conditioning, in which a mouse, head-fixed over a freely-moving wheel, must learn to close its eye in response to presentation of a small drifting grating stimulus (500 ms) prior to delivery of an air puff to the ipsilateral cornea (Fig. 1a, Extended Data Fig. 1, see methods)^25,26^. Simultaneous video recording of the eyelid position allowed us to determine correct (presence of conditioned response, CR) versus incorrect (presence of only unconditioned response, UR) performance on individual trials (Fig. 1a). Mice (n=39) readily learned this task over a few days with a consistently low false alarm rate (spontaneous blinks, Fig. 1c). Importantly, V1 lesion contralateral to the visual stimulus in naive mice (n=8) prevented learning (Fig. 1c), suggesting that the cortex is necessary for this behavior.

**Fig. 1:**
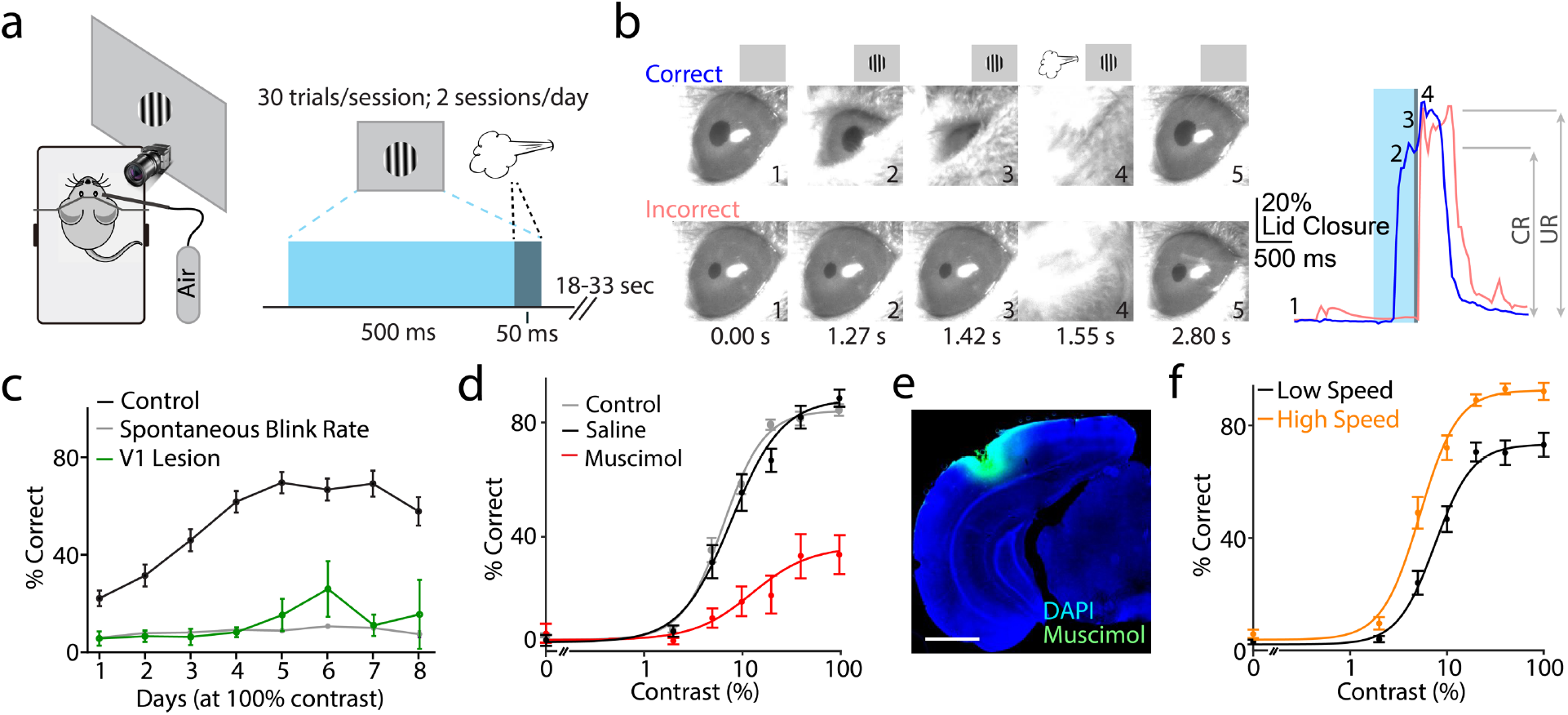
Visual cortex is necessary for visually cued eyeblink conditioning. **a**, Schematic illustration of the behavioral setup (left) and trial structure (right). **b**, Left, example video frames illustrating a correct conditioned blink (upper images) and incorrect unconditioned blink (lower images). Right, time course of lid closure for correct (blue) and incorrect (pink) trials shown in left panel, with timing of individual frames (numbers), visual stimulus (blue bar), and air puff (gray bar) indicated. **c**, Average ± SEM (n=39 mice) performance over learning for control (black) and V1 lesioned (green, n=8 mice) groups. Timing of the visual stimulus and air puff are shown above. Spontaneous blink rate per 450 ms period is shown (gray). **d**, Average ± SEM contrast-dependent performance for all control mice (gray, n=39) and mice injected with saline (black) and muscimol (red) into V1 on alternating days (n=11). R_Max_-saline 86.5%, R_Max_-muscimol 28.2%; p<0.0001, Permutation Test. **e**, Example showing extent of fluorescent muscimol infusion into V1. Scale bar is 1 mm. **f**, Average ± SEM (n=39 mice) performance separated into high (orange) and low (black) locomotion speed trials. R_Max_-high speed 88.7%, R_Max_-low speed 71.1%; p<0.0001, Permutation Test.

During training, the visual stimulus was set to 100% contrast. After mice demonstrated stable performance, the task was changed to present stimuli of varying contrast, allowing us to determine the average contrast-dependent behavior (Extended Data Fig. 1, Fig. 1d). Fitting these data with a hyperbolic ratio function (see methods) revealed the mean group performance (RMax: 81.2±1.9% correct, c50: 7.8±0.7% contrast, n=39 mice, Fig. 1d). In addition, both CR:UR ratio (response magnitude) and response latency were monotonically modulated by contrast (Extended Data Fig. 1). Unilateral inactivation of V1 using injection of the GABAergic agonist muscimol significantly impaired performance across contrasts (R_Max_-saline: 86.5%, R_Max_-muscimol: 28.2%, p<0.0001; Permutation Test, n=11 mice, Fig. 1d, e), further suggesting that V1 is necessary for visual perception in this task. Changes in behavioral state are also known to modulate cortical activity and perception^27–31^. We monitored locomotion and found that high speed trials exhibited significantly enhanced performance (R_Max_-High Speed: 88.7%, R_Max_-Low Speed: 71.1%, p<0.0001; Permutation Test, n=39 mice, Fig. 1f). Locomotion was also correlated with larger CR magnitude (Extended Data Fig. 1, Extended Data Table 1), consistent with state-dependent modulation of cerebellar motor control^32^. Notably, similar results were seen for pupil dilation (Extended Data Fig. 1, Extended Data Table 1). Overall, these results suggest that visually cued eyeblink conditioning provides a useful approach for investigating V1-dependent perceptual ability.

We next set out to determine how V1 neurons participate in visually cued eyeblink conditioning. Cortical outputs to the lateral pons may serve to direct sensory information to cerebellar structures critical for establishing associations between conditioned and unconditioned stimuli^25^. Previous work in cats suggested that corticopontine projections arise from multiple visual areas^33,34^. Using adenoassociated viral (AAV) tracing, we found that mouse V1 sends a strong projection to subcortical areas including the pons and dorsal striatum (Fig. 2a). Retrograde tracing from the pons labeled a subpopulation of layer 5 pyramidal neurons (PNs) in V1 as well as secondary visual areas and the retrosplenial cortex (Fig. 2b). Consistent with earlier work from our lab^17^, these corticopontine (CPn) cells were largely non-overlapping with co-mingled cells in layer 5 projecting to the ipsilateral dorsal striatum (CSt, Fig. 2b). As both corticostriatal and corticobulbar outputs have been linked to sensorimotor function^1,4^ and may extract distinct visual features from the environment^17^, we investigated whether these two populations might be differentially involved in the control of conditioned visuomotor behavior.

**Fig. 2:**
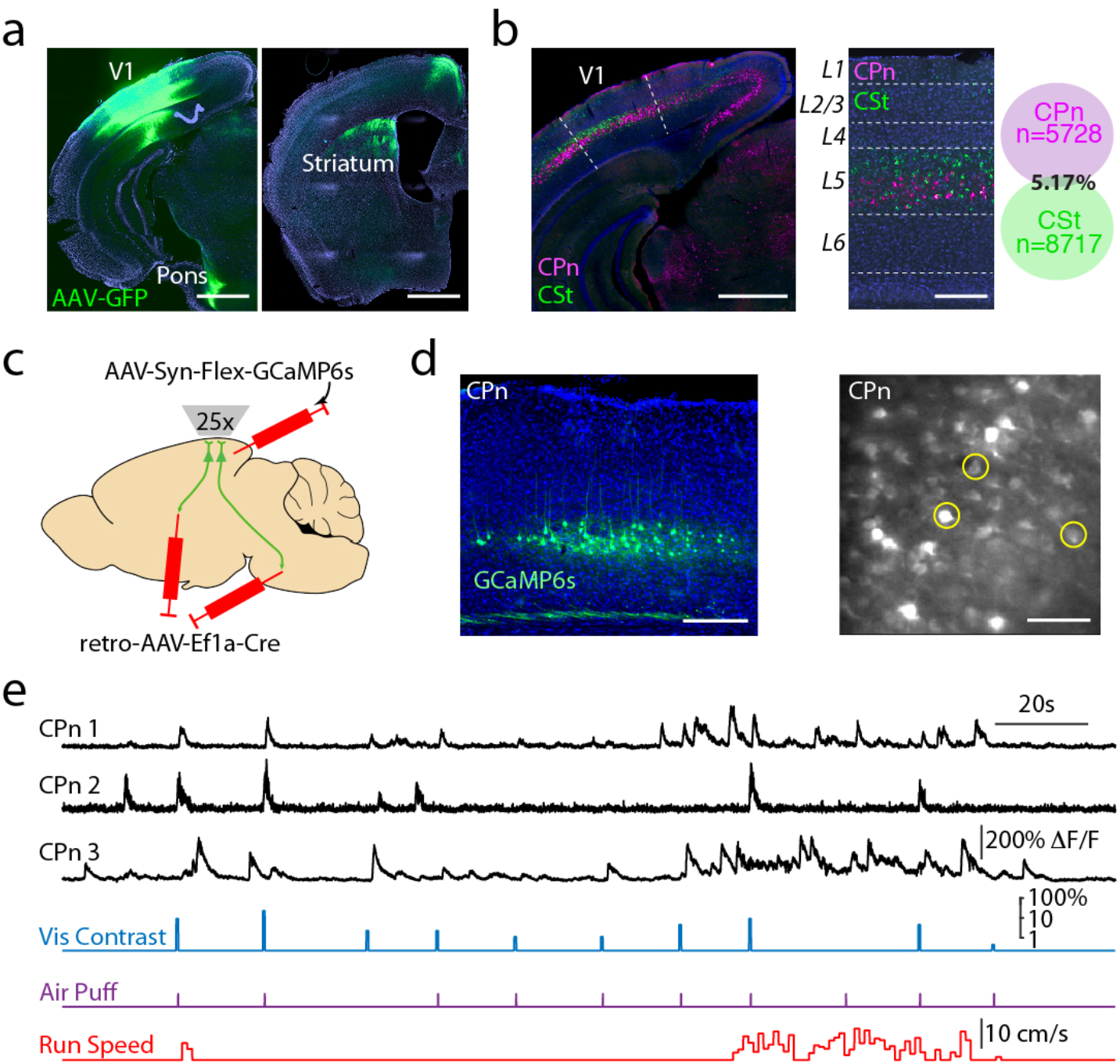
2-photon imaging of activity in projection-specific layer 5 subpopulations. **a**, Injection of AAV5-hSyn-GFP into V1 labels projections in both the ipsilateral pons (left) and ipsilateral dorsal striatum (right). Scale bars are 1 mm. **b**, Retrograde labeling of corticopontine (CPn, pink) and corticostriatal (CSt, green) cells with dual-color cholera toxin subunit B injections into ipsilateral pons or striatum reveals two non-overlapping populations that are mostly limited to layer 5 (left, center). Scale bars are 1 mm and 200 μm. Co-localization of CPn and CSt neurons (right, n=3 mice). **c**, Schematic illustrating intersectional viral approach for conditional expression of the calcium indicator GCaMP6s in layer 5 PNs projecting to either the ipsilateral dorsal striatum or pons. **d**, Left, example ex vivo image showing restricted expression of GCaMP6s in layer 5 corticopontine (CPn) neurons. Scale bar is 200 μm. Right, example in vivo 2-photon imaging of corticopontine neurons at 525 μm depth. Scale bar is 100 μm. **e**, Example traces showing simultaneous recordings of three CPn neurons from **d**, timing of visual stimuli and air puff, and continuous running speed.

To monitor the activity of different layer 5 PNs in task performance, we drove expression of Cre recombinase in either CPn or CSt cells using a retrograde AAV vector (retroAAV-Ef1a-Cre)^35^ injected into either the pons or striatum. We then drove conditional expression of the genetically encoded calcium indicator GCaMP6s^36^ with a second vector (AAV-Syn-Flex-GCaMP6s) injected into ipsilateral V1 (Fig. 2c). This approach resulted in selective labeling of the targeted subpopulation, compatible with monitoring neuronal activity in layer 5 of behaving mice via 2-photon microscopy through a chronically implanted cranial window (Fig. 2d). Importantly, we previously showed that GCaMP6s signaling of somatic spiking does not differ between subtypes of layer 5 PNs^17^.

Neuropil-subtracted fluorescence signals (ΔF/F) were collected and combined across 4-5 days during the varied contrast phase of training (Fig. 2e, Extended Data Fig. 2). Cells exhibited spontaneous and visually-evoked activity, as well as responses to the air puff and modulation correlated with locomotion (Fig. 2e). Consistent with previous reports in monkeys^37^, both populations also exhibited responses to spontaneous eyeblinks (Extended Data Fig. 3). To avoid contamination of the measured visual response, we quantified the evoked activity as the average signal in the 300 ms window following stimulus onset (prior to the blink response), subtracting the preceding 300 ms baseline period (Fig. 3a, Extended Data Fig. 3). Visually responsive neurons (see Methods) exhibited monotonically increasing output with increasing log-transformed visual contrast that was well-fit with a rectified linear contrast response function (CRF, Fig. 3a). This approach gave similar results to a sigmoid hyperbolic ratio function and yielded a single slope variable that could be compared across groups (Extended Data Fig. 4, Extended Data Table 1).

**Fig. 3:**
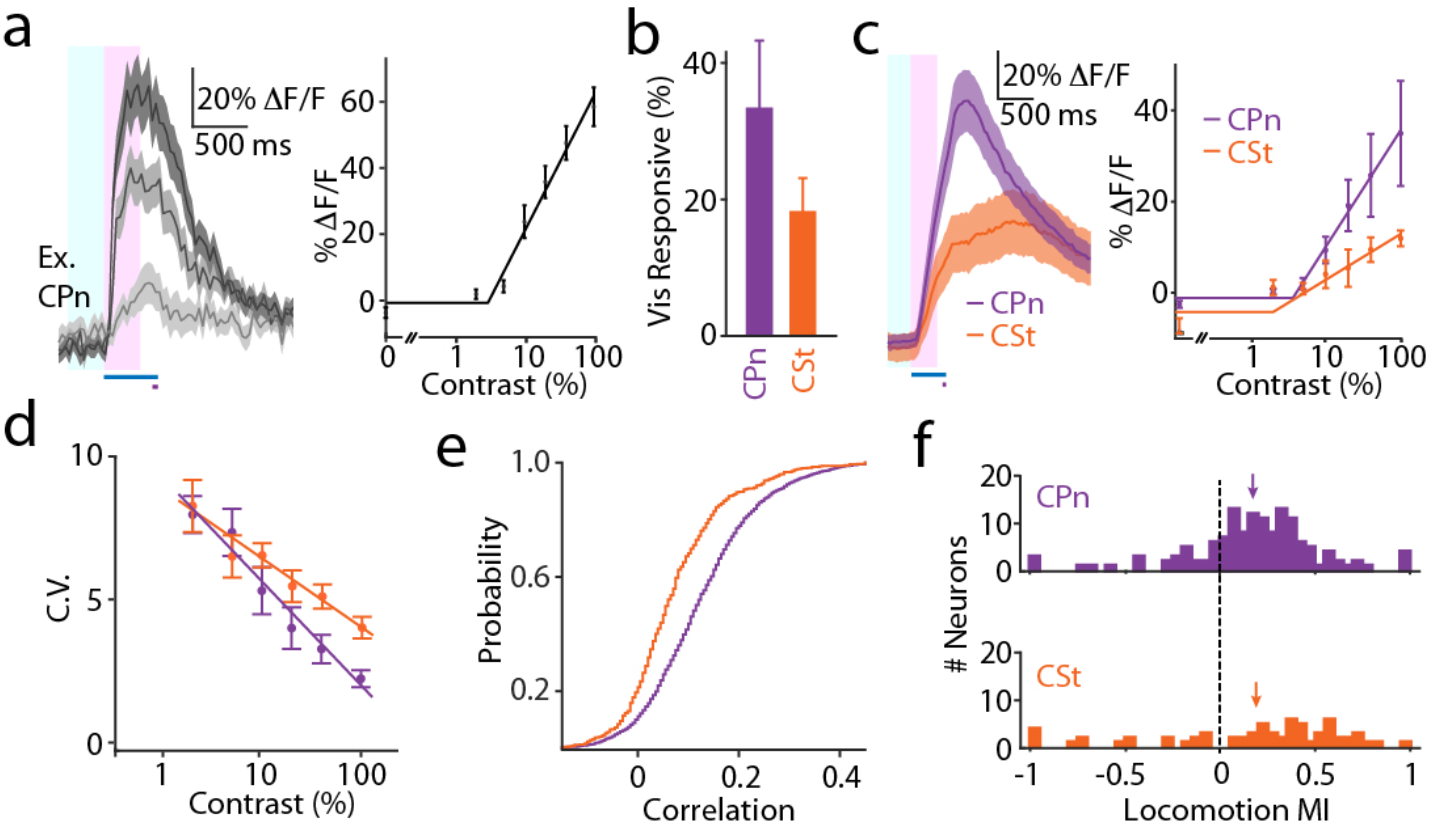
Visually evoked activity varies between layer 5 subpopulations. **a**, Left, average ± SEM visual responses from an example cell (CPn 1 from Fig. 2e) at 5%, 20%, and 100% contrast. Timing of visual stimulus (blue bar) and air puff (gray bar) are shown. Intervals for measuring baseline activity (light blue window) and visual response magnitude (pink window) are shown. Right, average ± SEM contrast-dependent response magnitudes and rectified linear curve fit for the cell. **b**, Average ± SEM proportion of visually responsive neurons in CPn (purple, n=6 mice, n=138 cells) and CSt (orange, n=6 mice, n=66 cells) cohorts. CPn 34.1±10.6%, CSt 18.4±5.2%; p=0.24, Mann-Whitney U Test. **c**, Left, population average ± SEM visual responses at 100% contrast for CPn (n=6 mice) and CSt (n=6 mice) cohorts. Right, population average ± SEM contrast-dependent response magnitudes for CPn and CSt cohorts. CPn slope 0.26±0.09, CSt slope 0.10±0.02; p=0.015, Mann-Whitney U Test. **d**, Population average ± SEM contrast-dependent coefficient of variation. CPn slope −3.40±0.42, CSt slope −2.02±0.28; p=0.04, Mann-Whitney U Test. **e**, Cumulative distributions of pairwise noise correlations for the two cohorts. CPn, 9453 pairs, median=0.126; CSt, 2145 pairs, median=0.049; p<0.0001, Kolmogorov-Smirnov Test. **f**, Histograms illustrating the distribution of locomotion modulation index values for the two cohorts. Average values indicated by arrows. CPn 0.18±0.03, n=138 cells, CSt 0.18±0.06, n=66 cells; p=0.93, t-test.

We found that a non-significantly larger fraction of CPn neurons was visually-responsive (34.1±10.6%, 138 neurons, 6 mice) versus CSt neurons (18.4±5.2%, 66 neurons, 6 mice, p=0.24, Mann-Whitney U Test, Fig. 3b). These CPn neurons exhibited significantly larger visual responses than CSt neurons (CPn slope: 0.26±0.09, CSt slope 0.10±0.02, p=0.015, Mann-Whitney U Test, Fig. 3c). CPn neurons also produced more reliable responses, measured as the slope of the contrast-modulated coefficient of variation (CPn slope: −3.4±0.42, n=6 mice; CSt slope: −2.02±0.28, n-=6 mice; p=0.04, Mann-Whitney U Test, Fig. 3d). Furthermore, consistent with our previous findings^17^, the functional connectivity of CPn neurons was greater than CSt neurons as measured by the pairwise noise correlations of visual responses (p<0.0001, Kolmogorov-Smirnov Test, Fig. 3e, Extended Data Table 1). Finally, both populations were similarly modulated by arousal, with cells in each group exhibiting positive or negative changes in visual responses associated with locomotion (Fig. 3f, Extended Data Table 1). These results suggest that, in mice trained on a conditioned visuomotor task, CPn neurons form a more robust and reliable network for distributing visual signals to subcortical structures.

To determine whether CPn and CSt cells carry information about task performance, we first separated visual responses into correct and incorrect trials to look at the average evoked activity. At contrasts near the behavioral threshold (10-20% contrast), CPn neurons demonstrated a larger average response in correct trials compared to incorrect trials, while CSt neurons exhibited similar responses for both trial outcomes (Fig. 4a-b). These results were significant across the range of contrasts (CPn slope_Correct_: 0.26, CPn slope_Incorrect_: 0.18, n= 6 mice, p=0.012, Permutation Test; CSt slope_Correct_: 0.11, CSt slope_Incorrect_: 0.11, n=6 mice, p=0.46, Permutation Test; Fig. 4c-d).

**Fig. 4:**
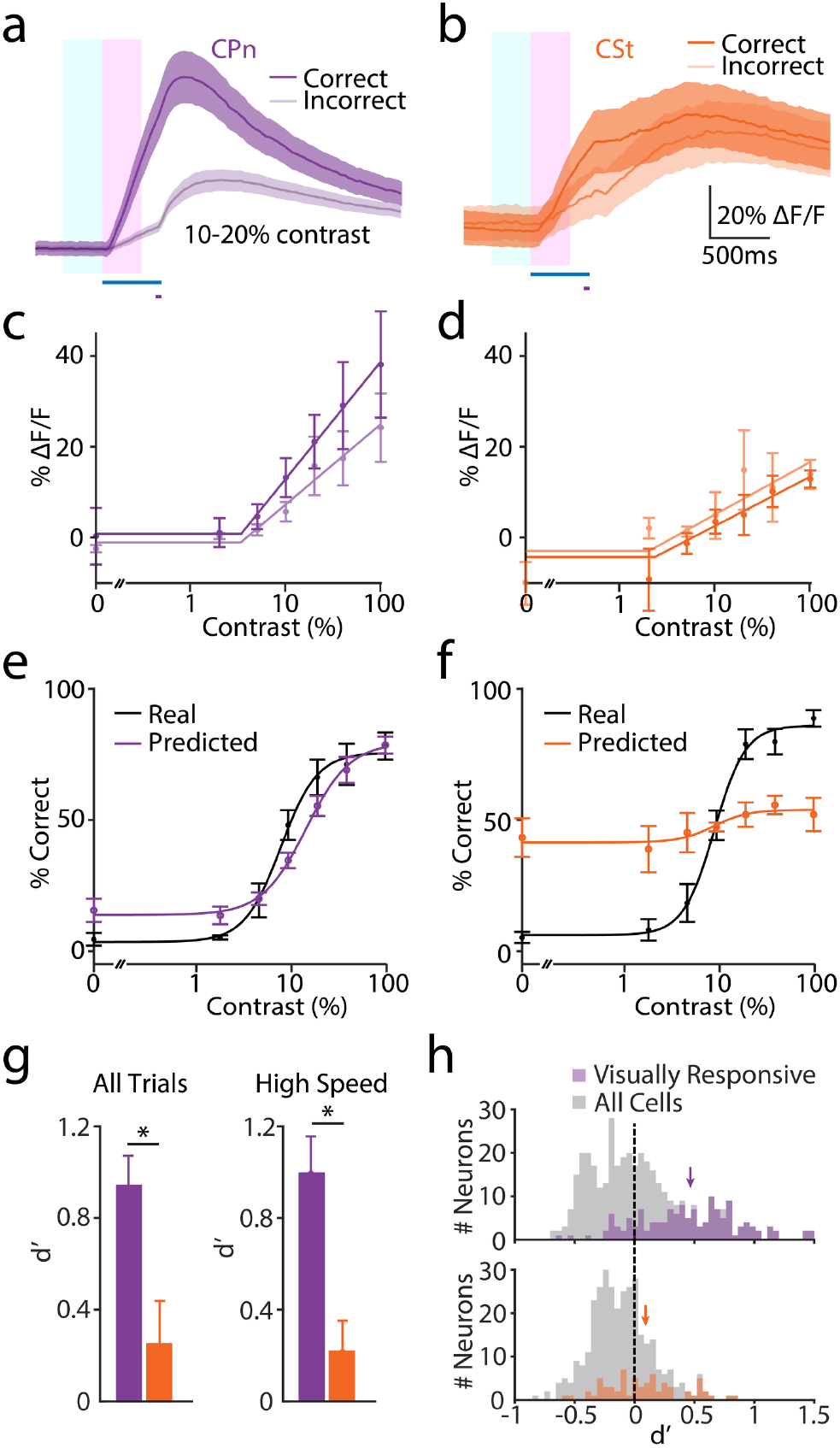
Behavior is selectively encoded by activity in CPn neurons. **a**, Average ± SEM (n=6 mice) visual responses at 10-20% contrast for CPn neurons separated by correct (dark) and incorrect (light) trials. Timing of visual stimulus (blue bar) and air puff (gray bar), and analysis windows (baseline, light blue; response, pink) are shown. **b**, As in a for CSt neurons (n=6 mice). **c**, Population average ± SEM contrast-dependent response magnitudes for CPn neurons (n= 6 mice) separated by correct (dark) and incorrect (light) trials. CPn Slope-Correct 0.26, Slope-Incorrect 0.18; p=0.012, Permutation Test. **d**, As in c for CSt neurons (n= 6 mice). CSt Slope-Correct 0.11, Slope-Incorrect 0.11; p=0.46, Permutation Test. e, Average ± SEM real (black) and predicted (based on CPn responses, purple) contrast-dependent behavior. **f**, As in e for CSt responses. **g**, Mean ± SEM discriminability (d’) for the ensemble decoders using CPn (purple, n= 6 mice) or CSt (orange, n= 6 mice) responses on all trials (left) or high locomotion speed trials (right). CPn all trials d’ 0.95±0.15, CSt all trials d’ 0.30±0.20; p=0.009, Mann-Whitney U Test. CPn high-speed trials d’ 1.00±0.16, CSt high-speed trials d’ 0.24±0.14; p=0.015, Mann-Whitney U Test. **h**, Histograms illustrating the distribution of d’ values for single visually-responsive CPn (purple) or CSt (orange) neurons, average values indicated by arrows. CPn average d’ 0.48±0.03, chance d’ −0.01±0.002, n=138 cells; p<0.0001, one-sided t-test. CSt average d’ 0.07±0.04, chance d’ −0.02±0.002, n=66 cells; p=0.033, one-sided t-test. CPn vs. CSt, p<0.0001, t-test. d’ values for all CPn (top) and CSt (bottom) neurons are shown in gray.

We then asked whether the output of the two populations could predict behavior on individual trials. We built a cross-validated logistic regression model using the activity of visually responsive CPn or CSt neurons to predict the outcome (blink or no blink) for each trial (see Methods). CPn activity reliably predicted psychometric behavioral data across all contrast levels (Fig. 4e), whereas CSt activity performed substantially poorer (Fig. 4f). We quantified behavioral discriminability (d’, see Methods) for the two populations and found that CPn output was significantly better than CSt output at predicting behavior (CPn d’: 0.95±0.15, CSt d’: 0.30±0.20, n= 6 mice per group, p=0.009, Mann-Whitney U Test, Fig. 4g). We obtained similar results when we limited the analysis to high running speed trials (Fig. 4g, Extended Data Table 1), suggesting that enhanced performance during locomotion is not directly linked to neuronal discriminability.

Behavior was encoded by the output of single neurons as well as by the population ensemble. For CPn cells, the average d’ value was 0.48±0.03 (n=138 cells, p<0.0001 vs. chance, one-sided t-test, Fig. 4h). For CSt cells, the average d’ value was 0.07±0.04 (n=66 cells, p=0.03 vs. chance, one-sided t-test; CPn vs. CSt, p=0.009, Mann-Whitney U Test, Fig. 4h, Extended Data Table 1). We also found that a small number of individual CPn neurons were capable of predicting behavior with similar accuracy as the population ensemble (Fig. 4h).

The preceding results indicate that significantly more information about individual trial performance is carried by CPn versus CSt neurons. We therefore wanted to determine whether the output of either population was necessary for normal task performance. We again used a retrograde AAV vector to express Cre recombinase in either of the two groups and a second AAV to conditionally express the optogenetic suppressor ArchT in V1 (Fig. 5a, b)^38^. This approach produced a similar density of infected cells (~7.5% of all layer 5 neurons) in each group (Extended Data Fig. 5, Extended Data Table 1). Mice were also implanted chronically with a guide cannula to enable placement of a fiber optic coupled to a 594 nm laser directly over V1 during the session.

**Fig. 5:**
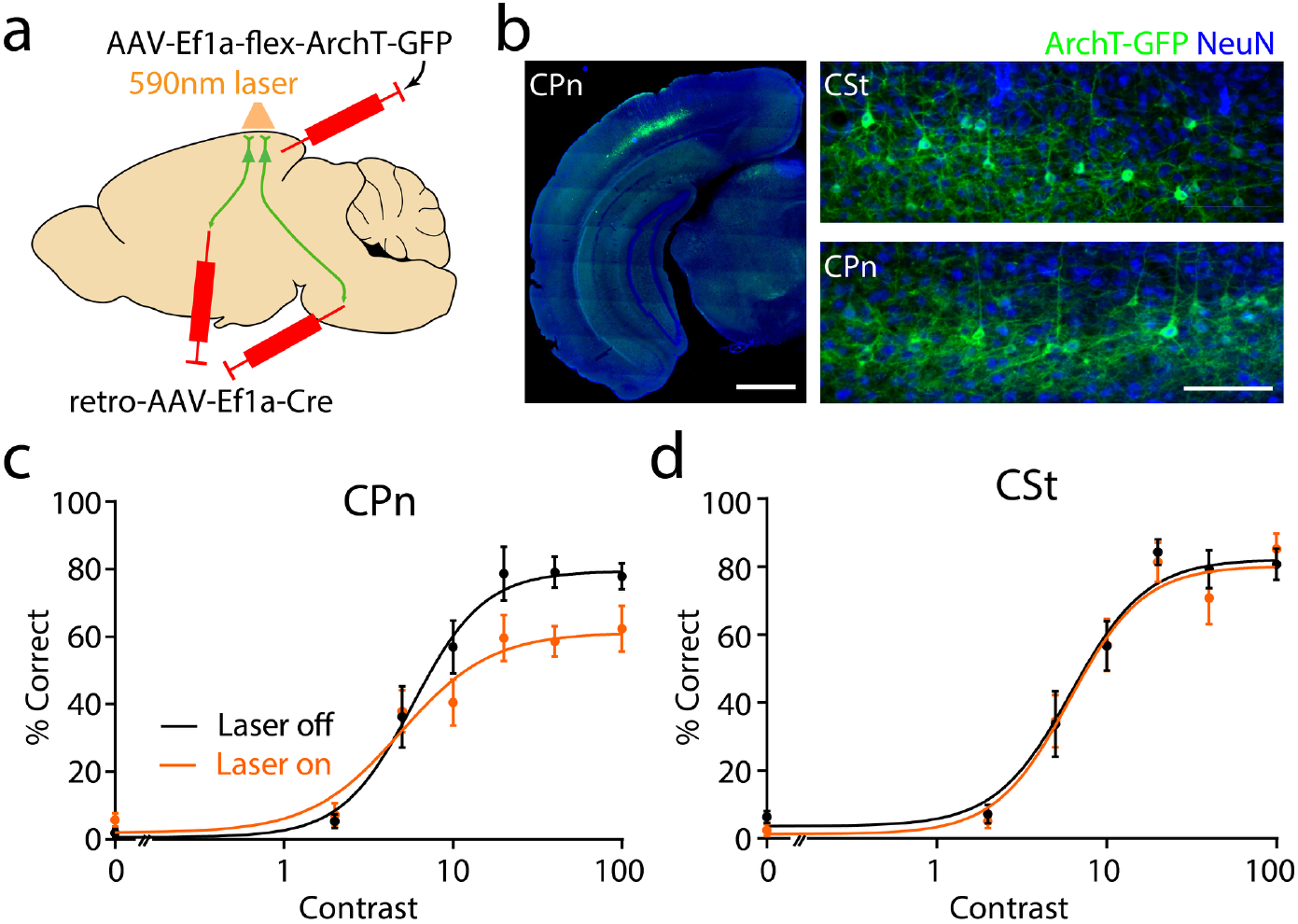
Optogenetic suppression of CPn activity disrupts behavioral performance. **a**, Schematic illustrating intersectional viral approach for conditional expression of ArchT-GFP in either CPn or CSt neurons. **b**, Left, example ex vivo image showing restricted expression of ArchT-GFP in V1 CPn neurons. Scale bar is 1 mm. Right, CPn and CSt neurons are similarly labeled. Scale bar is 100 μm. c, Average ± SEM (n=11 mice) performance for CPn cohort separated by trials with laser off (black) or on (orange). R_Max_ LaserOff 80.2%, R_Max_ LaserOn 60.5%; p<0.0001, Permutation Test. **d**, As in c for CSt cohort (n=12 mice). RMax LaserOff 79.3%, RMax LaserOn 77.7%; p=0.325, Permutation Test.

During the psychometric performance phase of training, a 3 second light pulse (120-160 mW/mm^2^ at the sample) was delivered to V1 on randomly interleaved trials, beginning 2 seconds before visual stimulus onset. Optogenetic inhibition of CPn neurons significantly suppressed behavior (RMax Laser_Off_: 80.2%, R_Max_ Laser_On_: 60.5%, n=11 mice, p<0.0001; Permutation Test, Fig. 5c). In contrast, inhibition of CSt neurons had no effect (R_Max_ Laser_Off_: 77.7%, R_Max_ Laser_On_: 79.3%, n=12 mice, p=0.24; Permutation Test, Fig. 5d). Importantly, optogenetic inhibition of either population did not directly interfere with motor ability, as unconditioned blink magnitudes were unaffected (Extended Data Fig. 5, Extended Data Table 1). These findings demonstrate that activity in CPn neurons not only encodes information about individual trial outcome but is required for optimal task performance.

## Discussion

In the present study, we developed a novel paradigm for probing the role of cortical microcircuits in visually-guided behavior. Our results indicate that mouse V1 is necessary for performing a visuomotor task that requires perception of a small contrast-modulated stimulus to produce a conditioned eyeblink response. Moreover, CPn (but not CSt) layer 5 PNs significantly encoded both sensory and behavioral information, and their activity was necessary for normal task performance. These findings are consistent with a model in which corticopontine projections provide a di-synaptic relay of visual signals to the cerebellum, where synaptic plasticity is thought to drive the association between conditioned and unconditioned stimuli^25,32,39^. Thus, cortical neurons are organized into physically interspersed but functionally distinct networks that can differentially participate in sensory-guided behavior.

Layer 5 PNs have long been recognized as a heterogeneous population based on morphology, electrophysiological properties, genetic markers, and projection targets^17,22–24^. Recent work from our lab and others suggests differential representation of visual stimuli, with brainstem-projecting PNs exhibiting broader orientation and spatial frequency tuning^17,40^. Synaptic connectivity of subgroups of layer 5 PNs with other PNs as well as with local interneurons may also differ, suggesting a basis for these distinct functional properties^11,15^. The possibility that different networks of cortical PNs, organized around common projection targets, might transmit distinct streams of behaviorally relevant information has been suggested for visual, somatosensory, and auditory areas^1,4,18,19^ Interestingly, modulation of corticostriatal activity was shown to alter behavioral output in an auditory discrimination task^4^, suggesting that layer 5 subnetworks may be differentially engaged by sensory inputs on the basis of current expectations or environmental demands. It will be very interesting to determine whether sensory representations in various cortical populations are plastic in response to passive experience versus task learning.

Recent studies have called into question the role of neocortex in learning and performance of basic sensory and motor tasks^3,41^. Here, we find that visually cued eyeblink conditioning is disrupted by V1 lesion, acute V1 inactivation, or selective optogenetic suppression of CPn cells, suggesting a necessary role for cortical activity. The reason for this discrepancy is unclear, though the aversive conditioning nature of our task and the specific involvement of brainstem-projecting PNs may distinguish our results from those obtained with appetitive, operant behavior. Indeed, our results, in combination with findings from other groups, indicate that caution is necessary when attempting to generalize conclusions across behavioral paradigms.

Our results are also consistent with studies demonstrating that behavioral state can modulate both perceptual ability, neuronal activity, and motor learning in mice^28,29,32^. The cellular mechanisms of for this effect remain unclear, with potential contributions from neuromodulatory systems^31,42,43^ and top-down feedback from other cortical regions and engagement of local inhibitory circuits^44,45^. Interestingly, our imaging data indicate that different layer 5 PNs exhibit either increases or decreases in activity during arousal without an overall change in discriminability, suggesting that the links between internal state, cortical signaling, and behavioral output are not straightforward.

In conclusion, our study demonstrates that layer 5 CPn neurons play a key role in the representation of visual information that is necessary for performing a conditioned motor behavior. This work supports the projection-specific parcellation of cortical projection neurons and provides strong evidence for the functional segregation of cortical microcircuits, even within a single layer. The formation and preservation of these distinct but physically interspersed networks are likely to be critical processes during cortical development, and loss of this specificity may ultimately contribute to abnormal sensorimotor behaviors associated with synaptic alterations in neuropsychiatric disorders.

## Acknowledgements

The authors wish to thank Dr. Daeyeol Lee for help with statistical analyses. We also thank Dr. Jessica A. Cardin and members of the Cardin and Higley laboratories for helpful discussions during the preparation of this manuscript.

## Author Contributions

LT and MJH designed the research and wrote the paper. LT performed all experiments and analyzed the data.

## Data Materials and Availability

All data are present in the paper and supplementary materials.

## Supplementary Materials

Materials and Methods

Table S1

Figures S1-S6

