## Extended Data for "Layer 5 circuits in V1 differentially control visuomotor behavior"

**This PDF file includes:**

Materials and Methods
Figures S1-S5
Table S1
Captions for Figures S1-S5

**Materials and Methods:**

Animals

All animal handling was performed in accordance with the Yale Institutional Animal Care and Use Committee and federal guidelines. Wild type C57/bl6 male mice were obtained from Envigo and group housed in standard cages on a 12h light-dark cycle.

Transcranial injection of retrograde tracer and virus

All surgical procedures were carried out in juvenile mice (P22-28 for viral injection, P30-40 for cholera toxin labeling) under isoflurane anesthesia. For retrograde labeling of layer 5 PNs, 100 nl of cholera toxin subunit B (CTB)-Alexa Fluor-555 or CTB-Alexa Fluor-488 (ThermoFisher) was injected via glass micropipette into the lateral pons (AP: -3.1, ML: 1.2, DV: 4.7 relative to Bregma) or dorsal striatum (AP: 0.5, ML: 1.7, DV: 1.8), respectively. To express either GCaMP6s or ArchT-GFP in specific layer 5 PN populations, 700 nl of retro-AAV-Ef1$\alpha$-Cre (Addgene) was injected into either the pons or striatum. A second 700 nl injection of AAV5-hSyn-Flex-GCaMP6s or AAV5-hSyn-Flex-ArchT-GFP, Addgene) was made into V1 (AP: -3.5, ML: 2.5, DV: 0.45).

Histology

Ten days after CTB injection or at the conclusion of behavioral experiments, mice under isoflurane anesthesia were perfused transcardially with phosphate buffer (PB) followed by 4% paraformaldehyde (PFA) in (PB). Brains were post-fixed in 4% PFA overnight at 4 °C. 40 $\mu$m-thick coronal sections containing the primary visual cortex were cut on a vibratome (Leica VT1000) and washed in PB.

For CTB-injected tissue, slices mounted on glass microscope slides and cover-slipped with DAPI-containing medium (Prolong, Invitrogen). For NeuN staining on ArchT-GFP-expressing tissue, slices were pretreated in blocking solution (2% BSA, 10% NGS, 0.5% Triton X100 in PB) for 4 hours followed by incubation in primary antibody (rabbit anti-mouse NeuN, Invitrogen) for 24 hours at 4 °C. Slices were then incubated in secondary antibody (goat anti-rabbit-Alexa Fluor 594, Invitrogen) for 2 hours at room temperature and mounted on glass slides.

Slices were imaged on an upright Olympus BX53 fluorescent microscope. A rectangular region of V1 layer 5 (600 µm in width, 350-550 µm in depth from pia) was cropped from the image of each section. For CTB retrograde tracing, a rectangular region of V1 layer 5 (600 $\mu$m width, 350-550 $\mu$m depth from pial surface) was collected. Labeled cells in each channel were identified and counted using ImageJ. For estimating ArchT-GFP-expression, we used CellProfiler to expand NeuN-identified nuclei by 2 pixels to estimate somatic area. Neurons were classified as ArchT-GFP-positive if >60% of soma pixels displayed GFP fluorescence.

Cannulation, head-posting, and window implantation

Mice were anesthetized with isoflurane, the scalp was resected, and a small craniotomy was made over V1. For acute inactivation experiments, a guide cannula (26 gauge, Plastics One) was lowered to the brain surface. For optogenetic experiments, a larger guide cannula (21 gauge, Plastics One) was used. The cannula was secured via two screws set into the skull and dental cement (Metabond, Parkell). For lesion experiments, a similar procedure was used, and V1 was surgically resected using a small metal probe. A custom titanium head-post was then fixed to the skull using dental cement. A blocking stylus was placed in the cannula to prevent contamination. For imaging experiments, a ~4 mm square craniotomy was made over left V1. A bilayer imaging window consisting of a 5x5mm glass coverslip bonded to a 3.5x3.5 mm inner glass using ultraviolet-curing adhesive (Norland Products) was inserted in the craniotomy and secured to the skull with Metabond.

Visual stimulation

Sinusoidal drifting gratings were generated using Psychtoolbox-3 in MATLAB and presented on a gamma-calibrated LCD monitor (17 inches) with a spatial resolution of 1280x960, frame rate of 60 Hz, and mean luminance of 45 cd/m$^2$. Unilateral stimuli had a 20° visual angle, temporal frequency of 2 Hz, spatial frequency of 0.04 cycles per degree, and orientation of 180°. During the training phase, visual stimuli were set to 100% contrast. During psychometric testing, contrast varied (0%, 2%, 5%, 10%, 20%, 40%, 100%) pseudorandomly. For non-imaging experiments, stimuli were presented in the center of the LCD screen. For calcium imaging experiments, the stimulus was fixed in one of nine 3x3 sub-regions that evoked the largest population response in the field of view.

Behavioral setup

The mouse was head-fixed on a freely-moving wheel (15 cm diameter) in a darkened and sound-attenuating chamber. The LCD screen was positioned on the right side at a distance of 22 cm and normal to the right eye. Timing of visual stimuli was recorded using a photo-diode that detected a small luminance signal in the corner of the screen. Brief air puffs (10-12 psi) were generated with a compressed air tank coupled to a solenoid (Clark Solutions). The air puff was directed to the right cornea using a 14-gauge stainless steel cannula positioned ~5 mm from the eye. Timing of the air puff was coordinated with the visual stimulus using custom-written MATLAB codes through a NI-DAQmx board (PCIe-6315, National Instruments) at a sampling rate of 5 kHz. To monitor locomotion, a magnetic angle sensor (Digikey) was attached to the shaft of the wheel. Eyelid closure and pupil diameter were continuously recorded using a monochromatic CMOS camera (PointGrey FlyCapture3) at a frame rate of 33 fps. An infrared LED array was directed toward the animal's to illuminate the eye. All signals, including the timing of the visual stimuli, the air puffs, the wheel position, and video frame ticks were digitized (5 kHz) and recorded through the NI-DAQmx board for non-imaging experiments. For calcium imaging experiments, all signals as well as the microscope resonant scanner frame ticks were digitized (5 kHz) and collected through a Power 1401 (CED) acquisition board using Spike 2 software.

Muscimol inactivation

Saline solution consisted of artificial cerebrospinal fluid (ACSF) adjusted to pH of 7.4. Muscimol (1 μg/μl, Sigma) or bodipy-tagged muscimol (1μg/μl, Thermo-Fisher, final day only) was dissolved in ACSF. At the conclusion of the final behavioral day, mice were perfused and their brains sectioned to validate the infusion site and extent. On alternating days, either saline or muscimol-containing saline was injected through the implanted cannula (0.5 µl at a speed of 0.25 µl/min). The cannula was capped with a blocking stylus, and the mouse was returned to its home cage for 15 min prior to behavioral testing.

Induced pupil dilation

The mouse was lightly anesthetized using isoflurane, and a drop of saline (0.9%) or atropine solution (1% atropine in saline) was applied to the right eye. The mouse was allowed to recover in its home cage for 30 minutes prior to behavioral testing.

Optogenetic suppression of cortical activity

Optogenetic experiments were carried out approximately 21 days following virus injection. A multi-mode optical fiber (400 µm diameter, 0.22 NA, ThorLabs) was coupled to a 594 nm diode-pumped solid state laser (Excelsior, Spectra Physics) whose output was first directed through a Pockels Cell (Conoptics) for power adjustment. Power was adjusted to 120-160 mW/mm$^2$ at the output end of the fiber, checked at the start of each session. The fiber was inserted into the guide cannula so that the tip was just above the pial surface. In half of the trials (set pseudorandomly), the light was turned on 2 seconds prior to the visual stimulus onset and turned off after 3 seconds via a electronically gated shutter (Uniblitz).

Behavioral Training

For all behavioral experiments, mice were habituated to being head-fixed on the wheel for at least three days prior to the start of training. Each trial started with the onset of the 500 ms visual stimulus (CS). The 50 ms air puff (US) was delivered at 450 ms to co-terminate with the CS. For non-imaging experiments, each training consisted of 60 CS-US pairings 100% contrast. After demonstrating stable conditioned responses (>50% correct) for 2-3 consecutive days, mice were moved to the performance phase and 60 CS-US pairings with varying contrasts were presented. Mice that did not learn after 14 days of training (n=7/46) were excluded from further experiments. For calcium imaging experiments, performance sessions also included 12 CS-only trials at varying contrasts and 6 US-only trials (0% contrast). For all sessions, trials were separated by an exponentially-distributed inter-stimulus-interval (ISI) ranging from 18 to 33 seconds.

Calcium imaging

Imaging experiments were conducted approximately 21 days after virus injection and 12 days after surgical implantation. Imaging was carried out using a resonant scanner-based two-photon microscope (MOM, Sutter Instruments) through a 25x, 1.05 NA objective (Olympus) coupled to a Ti:Sapphire laser (MaiTai DeepSee, Spectra Physics) tuned to 920 nm for GCaMP6s. Emitted light was collected through gallium arsenide phosphide photomultiplier tubes (Hamamatsu). To prevent light contamination from the display monitor, the microscope was enclosed in blackout material that extended to the head-post. Images were acquired using ScanImage 2017 (Vidrio)(*1*) at ~30 Hz and a resolution of 256x256 pixels (290x290 μm). Layer 5 PN somata were imaged at ~450-600 μm depth relative to the brain surface, as previously published (*2*). The same field of view was imaged on each day and identified neurons were tracked across sessions.

Behavioral analysis and statistics

Eyeblink and pupil videos were analyzed offline with custom MATLAB scripts. To extract eyelid closure, gray-scale images from a single session were binarized to maximize the contrast between the eye (white) and surrounding fur (black). A region of interest around the eye was manually defined, and the time-varying proportion of white pixels was used as a readout for eye closure. These data were normalized by the 5th and 95th percentile values for each session, resulting in a range of 0 to 1, corresponding to a fully open and fully closed eye, respectively. For each trial, the conditioned response (CR) was defined as the maximum eye closure within 450 ms of the visual stimulus onset (prior to air puff). The unconditioned response (UR) was defined as the maximum eye closure within a 500 ms window from the onset of the air puff. Trials were identified as correct if the CR:UR ratio was larger than 10%. CR:UR values were only analyzed for correct trials. Trials were excluded from analysis if the eye closed >10% within a 2 second window prior to visual stimulus onset. Spontaneous blinks (>10% full eye closure) were detected during the inter-stimulus-intervals. Spontaneous blink rate was calculated as the average number of blinks per 450 ms interval to compare with the behavioral analysis window.

To extract pupil size, gray-scale images were binarized (independently from analyses of eyeblink) to maximize the contrast between the pupil (black) and surrounding sclera (white). Pupil size was determined as the time-varying proportion of black pixels in a region of interest covering the eye. Pupil data were normalized by the 5th and 95th percentile values for each session. For each trial, arousal state was determined as the average pupil size during the 2 seconds prior to visual stimulus onset. Trials were classified into large and small pupil classes using the median value across mice.

To determine running speed, the rotational wheel position data during each session were binned over 100 ms intervals and transformed into velocity. For each trial, average running speed during 2 seconds prior to visual stimulus onset was measured and classified into high and low classes using a threshold of 1 cm/s.

All trials across days from the performance phase (using varied contrast) were concatenated for analysis. Group performance was calculated by averaging the % correct at each contrast across animals and curve fitting these values with a hyperbolic ratio function:

$$\%\,Correct = baseline + \frac{R_{max} \times Contrast^{Power}}{Contrast^{Power} + {c_{50}}^{Power}}$$

For fitting purposes, baseline was the spontaneous blink rate at 0% contrast, c50 was restricted to <50% contrast, and the power was restricted to between 1 and 3. In the present work, we limited statistical analyses of behavioral performance to the calculated $R_{Max}$ value. Contrast dependence of both the CR:UR ratio and the RT were fit with linear functions for contrast values above 2%.

To address whether behavioral variables (e.g., $R_{Max}$, CR:UR ratio, RT) varied by muscimol treatment, arousal, or optogenetic manipulation, we then conducted a stratified permutation analysis to maximize the use of data points across contrast levels and allow pairwise comparisons between conditions within each mouse. Trials within mouse at a single contrast level were regarded as a stratum. For each permutation, condition labels (e.g., muscimol versus saline) were randomly permuted (shuffled) within each strata so that the number of trials per condition per contrast per animal remained unchanged. The performance at various contrast levels across all mice was then averaged using the permuted data and the population averages were then fit with the appropriate curve (hyperbolic ratio or linear) for the two paired conditions. We then calculated the ratio of the parameter of interest (e.g., $R_{Max}$) from these curves. This process was repeated 10,000 times to generate a null distribution for the group average parameter. The p value for each parameter was calculated by summing the proportion of points beyond the actual value in the null distribution.

This permutation approach maximizes power but requires curve fitting of data and does not allow for a simple, paired statistical comparison. Therefore, to complement our analyses we carried out a direct pairwise test of parameter values across mice using data from trials at 100% contrast. The statistical results did not differ in any case between the two methods. Permutation results are presented in the Main text, while all results (Permutation and Pairwise) are provided in Supplemental Table 1.

Imaging analysis and statistics

Analysis was performed using custom-written scripts in MATLAB. $Ca^{2+}$ imaging data was motion-corrected using the Moco plugin for ImageJ (*3*), taking the first 200 frames of each movie as the template. Videos from successive days were translated onto the first-day template. Regions of interest were selected as previously described (*4*). Fluorescence (F) over time was measured by averaging all pixels in a given ROI, and the contamination from the surrounding neuropil was removed as previously described with a discounting coefficient of 0.70 (*2, 4*). $\Delta F/F$ was calculated as $(F-F_0)/F_0$, where $F_0$ was the lowest 10% of values from the neuropil-subtracted trace for each session.

The visual response of a neuron on a given trial was defined as the mean $\Delta F/F$ in a 300ms window after the visual stimulus onset, subtracting the mean $\Delta F/F$ over the 300 ms preceding the stimulus. A neuron was classified as visually responsive if its response at 100% contrast was significantly larger than those at 0% contrast using a one-sided Student's t-test. Only visually-responsive neurons were included in the analyses unless otherwise noted.

The contrast response function (CRF) of each visually responsive neuron was estimated using a 3-parameter rectified linear unit function (ReLU), as it yielded the largest R-square and the smallest adjusted root-mean-squared-error compared to sigmoidal fit (Supplemental Fig. 6):

$$\mathbf{\Delta F/F} = \textit{offset} + \log_{10}(\mathbf{Contrast}) \times \textit{Slope} \times \mathrm{I}\,(\mathbf{Contrast} > \textit{Threshold})$$

where offset is the spontaneous calcium activity at 0% contrast and slope indicates how fast the visual response increases with contrast, only when contrast is above threshold. The average visual response amplitudes, per contrast, of all the visually responsive neurons in each animal were grouped to generate a within animal CRF, fitted as described above. The coefficient of variation (standard deviation divided by mean) versus contrast was plotted similarly. Pairwise noise correlations between visually responsive neurons were calculated after Z-scoring response amplitudes across trials and concatenating across all trials. Noise correlations were defined as the Pearson's r-value across trials for each pair of cells. A modulation index for locomotion was calculated for each neuron as:

$$\mathrm{MI} = (\mathrm{Slope}_{\mathrm{High}}-\mathrm{Slope}_{\mathrm{Low}})/(\mathrm{Slope}_{\mathrm{High}}+\mathrm{Slope}_{\mathrm{Low}})$$

For calcium imaging experiments, data from all neurons within an animal were pooled for calculation of across-animal statistics. This approach is more conservative than an across-cells comparison which violates independency assumptions for most test. As the animal numbers were small (n=6 for CPn neurons and n=6 for CSt neurons), we used non-parametric analyses, specifically Mann-Whitney U Tests for non-paired samples and Wilcoxon tests for paired samples.

To quantify the relationship of neuronal activity to behavior, trials were classified into correct and incorrect, and analyzed via permutation, as described above. The trial-by-trial decoder was built to predict the behavioral performance of individual animals using visually evoked activity via simple logistic regression as follows. Let $x_i$ be a vector of visual responses for each trial for the i-th neuron, p is the probability of a correct trial, and m is the total number of recorded neurons in a mouse:

$$log\,\frac{p}{1-p} = \beta_0 + \sum_{i=1}^{m} \beta_i x_i + \varepsilon$$

Coefficients were estimated using maximum likelihood from 90% of trials and used to predict the 10% of the trials (10-fold cross-validation). A predicted trial was regarded as correct for p >=0.5. The predicted psychometric performance was then plotted against contrast using the actual contrast labels. To account for unbalanced numbers of neurons in the two subpopulations, each mouse in the CPn cohort was matched with a mouse in the CSt cohort and an equally sized subset of neurons was randomly selected for each pair. For each mouse, this selection process was repeated 15,000 times or the maximum combination of neurons, whichever was smaller.

The performance of each population to predict behavior was measured as a d' value, defined as: Z(model hit rate)-Z(model false alarm rate), where the function Z(p) with $p \in [0,1]$ is the inverse cumulative distribution function of a standard Gaussian distribution. A similar approach was used for constructing a single neuron decoder. A two-sample t-test was used to compare the cell-wise performance between subpopulations.

1. T. A. Pologruto, B. L. Sabatini, K. Svoboda, ScanImage: flexible software for operating laser scanning microscopes. *Biomed Eng Online* **2**, 13 (2003).
2. G. Lur, M. A. Vinck, L. Tang, J. A. Cardin, M. J. Higley, Projection-Specific Visual Feature Encoding by Layer 5 Cortical Subnetworks. *Cell Rep* **14**, 2538-2545 (2016).
3. A. Dubbs, J. Guevara, R. Yuste, moco: Fast Motion Correction for Calcium Imaging. *Front Neuroinform* **10**, 6 (2016).
4. T. W. Chen *et al.*, Ultrasensitive fluorescent proteins for imaging neuronal activity. *Nature* **499**, 295-300 (2013).

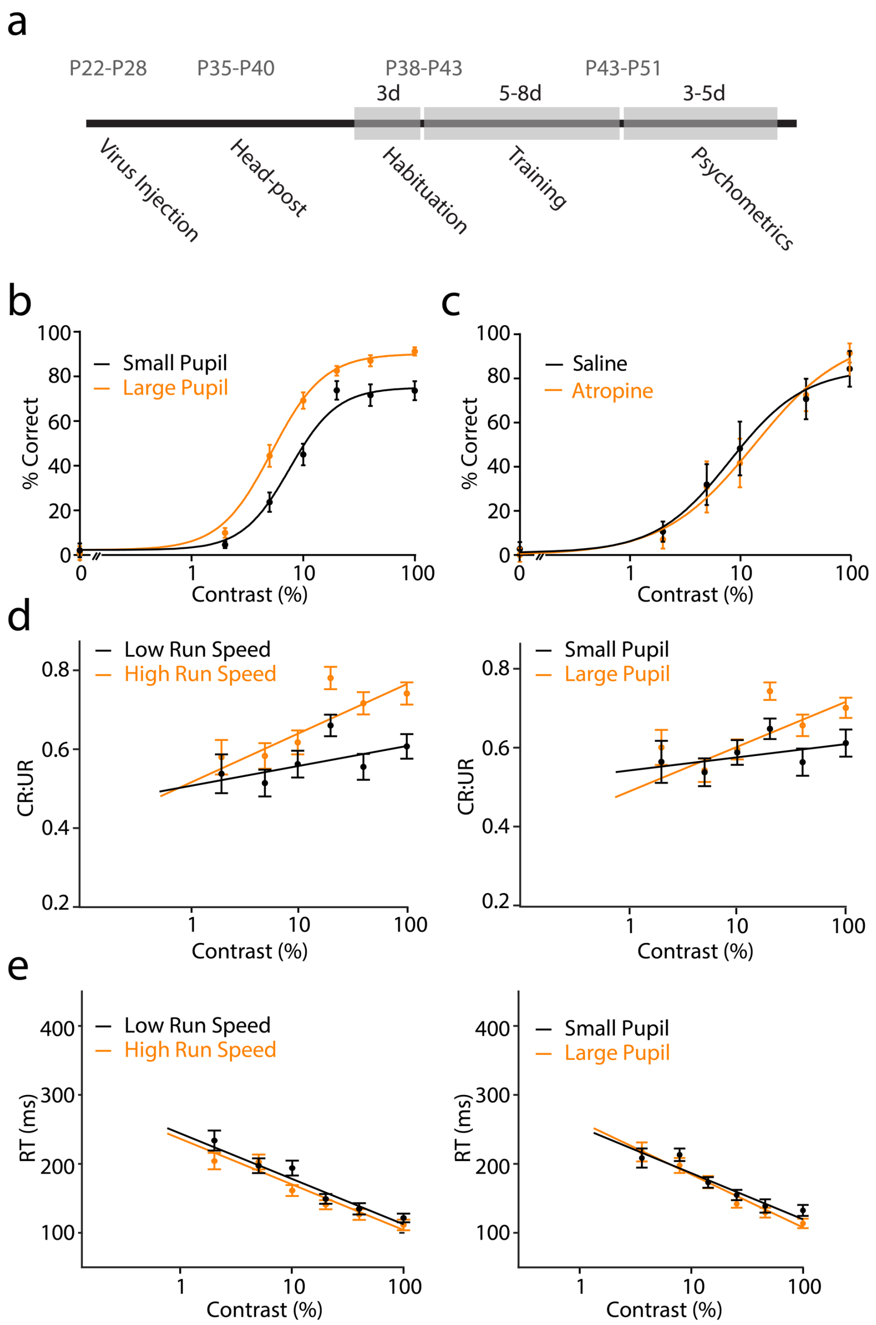


**Extended Data Fig. 1: Behavioral state is correlated with visual task performance.**
**a**, Timeline of experimental procedures. **b**, Average ± SEM (n=39 mice) performance separated into large (orange) and small (black) pupil trials. $R_{Max}$-Large Pupil 85.5%, $R_{Max}$-Small Pupil 69.9%; p<0.0001, Permutation Test. **c**, Average ± SEM (n=9 mice) performance measured 30 minutes after application of saline (black) or atropine (orange) to the eye. $R_{Max}$-Saline 72.4%, $R_{Max}$-Atropine 76.0%; p=0.365, Permutation Test. **d**, Left, average ± SEM (n=39 mice) relative amplitude of conditioned blinks (CR:UR) separated into high (orange) and low (black) locomotion speed trials. Slope-High Speed 0.126, Slope-Low Speed 0.055; p=0.0063, Permutation Test. Right, as in left panel for large and small pupil diameter. Slope-Large Pupil 0.115, Slope-Small Pupil 0.038; p=0.0034, Permutation Test. **e**, Left, average ± SEM response time (RT) separated by high and low running speed. Slope-High Speed -0.067, Slope-Low Speed -0.065; p=0.338, Permutation Test. Right, as in left panel for large and small pupil diameter. Slope-Large Pupil -0.066, Slope-Small Pupil -0.060; p=0.290, Permutation Test.

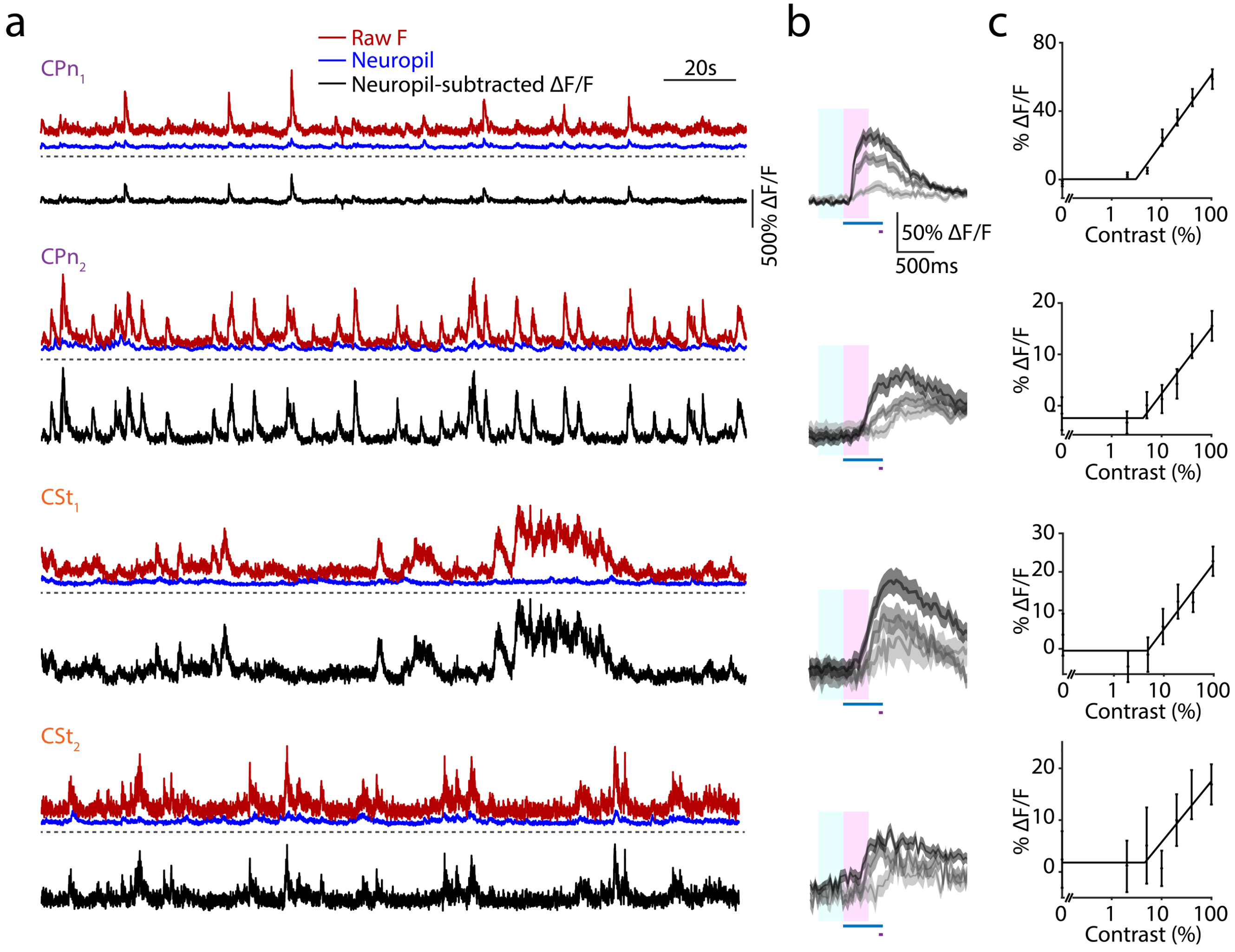


**Extended Data Fig. 2: Additional example calcium imaging traces and visual responses from CPn and CSt neurons.**
**a**, Examples of two CPn and two CSt neurons. Raw fluorescence (red), neuropil fluorescence (blue), and neuropil-subtracted ΔF/F (black) are shown for each. Dashed line indicates zero raw fluorescence. **b**, Average ± SEM visual responses at 5%, 20%, and 100% contrast (light, medium, dark gray) for each cell in **a**. Timing of visual stimulation (blue bar), air puff (gray bar), baseline analysis period (light blue window), and response measurement period (pink window) are indicated. **c**, Average ± SEM contrast-dependent response magnitudes for the example neurons, fit with a rectified linear curve.

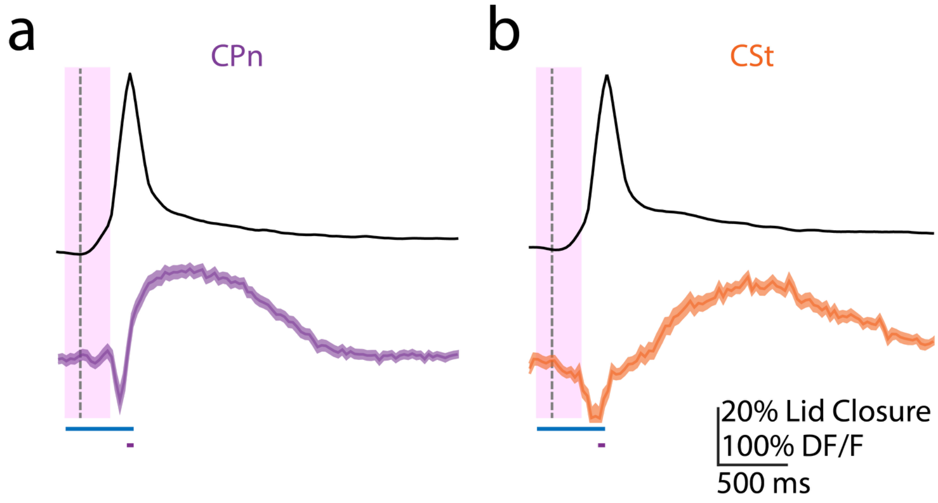


**Extended Data Fig. 3: Neuronal calcium transients in response to spontaneous blinks.**

**a**, Average lid closure for spontaneously detected blinks (black) and average ± SEM calcium transient (purple) for all visually-responsive CPn neurons. Dashed line indicates blink onset. Timing of visual stimulus (blue bar), air puff (gray bar), and response measurement (pink window) are displayed relative to the average onset of high contrast stimulus-evoked blinks (100 ms). Note the measurement window does not include contamination from the blink-evoked signal. **b**, As in **a** for visually-responsive CSt neurons.

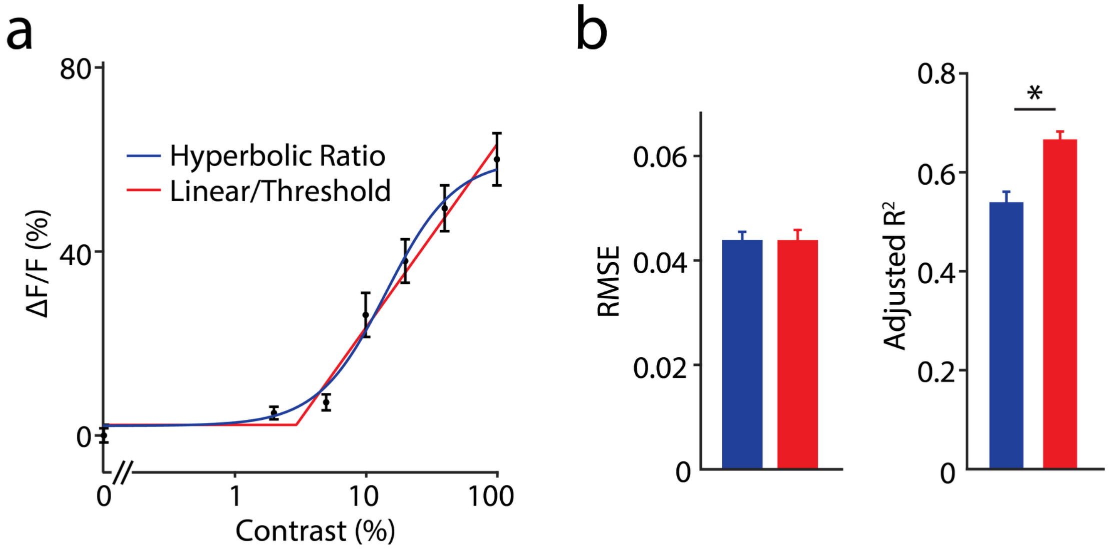


**Extended Data Fig. 4: Comparison of hyperbolic ratio and rectified linear curve fits for neuronal contrast sensitivity.**
**a**, Example, average ± SEM visual responses for a single neuron across multiple contrasts. Data are fit with either a hyperbolic ratio function (blue) or rectified linear function (red). **b**, Left, average ± SEM (n=204 cells) root-mean-squared error (RMSE) for the two fitting functions, n=204 cells. Hyperbolic Ratio 0.043±0.002, Linear/Threshold 0.043±0.002; p=0.498, Paired t-test. Right, as in left panel for adjusted $R^2$. Hyperbolic Ratio 0.54±0.02, Linear/Threshold 0.67±0.02, n=204 cells; p<0.0001, Paired t-test.

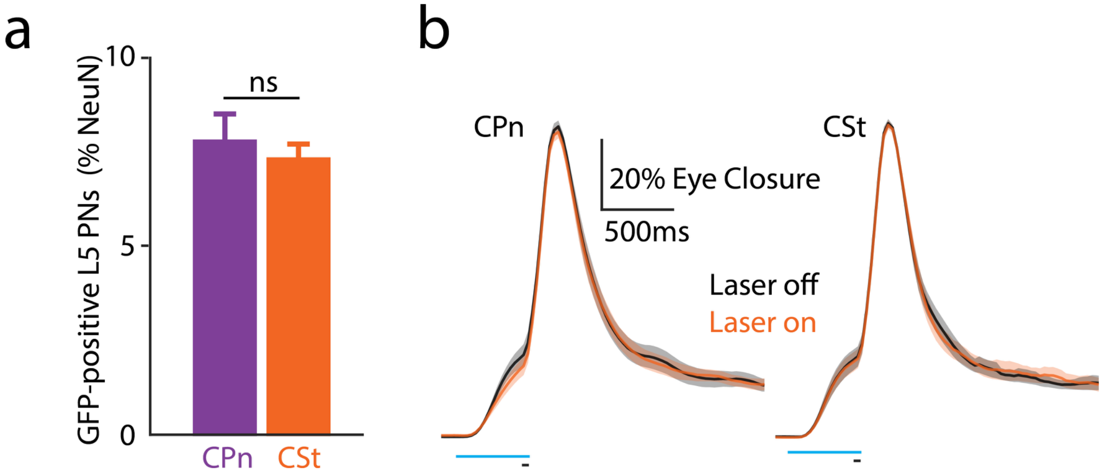


**Extended Data Fig. 5: Neuronal density of ArchT-GFP expression for CPn and CSt cohorts.**
**a**, Average ± SEM proportion of NeuN-identified layer 5 neurons expressing ArchT-GFP in either CPn (purple) or CSt (orange) cohorts. CPn proportion 7.76±0.75%, n=6 mice; CSt proportion 7.28±0.42%, n=6 mice; p=0.589, Mann-Whitney U test. **b**, Average ± SEM (n=11 and 12 mice, respectively) lid closure traces for all contrasts, separated by laser-off (black) or laser-on (orange) trials, for CPn (left) and CSt (right) cohorts. CPn UR amplitude-LaserOn 0.91±0.02, UR amplitude-LaserOff 0.89±0.02; p=0.178, Paired t-test. CSt UR amplitude-LaserOn 0.91±0.01, UR amplitude-LaserOff 0.92±0.01; p=0.748, Paired t-test.

**Table S1. Summary of all statistical analyses.**

| Figure | Comparison | Test | Statistic A | Value A | Statistic B | Value B | Test statistic | 95% Confidence Interval | p-value |
|---|---|---|---|---|---|---|---|---|---|
| **Behavior** | | | | | | | | | |
| Fig.1d | Muscimol vs. Saline performance (n=11 mice) | Permutation | Muscimol Rmax | 28.2% | Saline Rmax | 86.5% | A/B=0.33 | [0.83,1.19] | <0.0001 |
| | | Paired t-test | Muscimol 100% contrast | 28.9±5.4% | Saline 100% contrast | 89.3±2.9% | | | <0.0001 |
| Fig.1f | High-speed vs. Low-speed performance (n=39 mice) | Permutation | High speed Rmax | 88.7% | Low speed Rmax | 71.1% | A/B=1.24 | [0.94,1.10] | <0.0001 |
| | | Paired t-test | High speed 100% contrast | 92.1±3.1% | Low speed at 100% contrast | 73.1±4.2% | | | 0.0013 |
| Fig.S1b | Large pupil vs. Small pupil performance (n=39 mice) | Permutation | Large pupil Rmax | 85.5% | Small pupil Rmax | 69.9% | A/B=1.10 | [0.94,1.06] | <0.0001 |
| | | Paired t-test | Large pupil 100% contrast | 91.0±1.9% | Small pupil 100% contrast | 72.1±4.4% | | | 0.00011 |
| Fig.S1c | Atropine vs. Saline performance (n=9 mice) | Permutation | Atropine Rmax | 76.0% | Saline Rmax | 72.4% | A/B=1.05 | [0.79,1.27] | 0.365 |
| | | Paired t-test | Atropine at 100% contrast | 76.0±7.6% | Saline at 100% contrast | 74.2±8.1% | | | 0.820 |
| Fig.S1di | High speed vs. Low speed CR:UR (n=39 mice) | Permutation | High speed Slope | 0.126 | Low speed Slope | 0.055 | A/B=2.31 | [0.54,1.94] | 0.0063 |
| | | Paired t-test | High speed 100% contrast | 0.741±0.028 | Low speed 100% contrast | 0.607±0.031 | | | <0.0001 |
| Fig.S1dii | Large pupil vs. Small pupil CR:UR (n=39 mice) | Permutation | Large pupil Slope | 0.115 | Small pupil Slope | 0.038 | A/B=3.03 | [0.54,2.11] | 0.0034 |
| | | Paired t-test | Large pupil 100% contrast | 0.704±0.026 | Small pupil 100% contrast | 0.614±0.034 | | | 0.0012 |
| Fig.S1ei | High speed vs. Low speed RT (n=39 mice) | Permutation | High speed Slope | -0.067 | Low speed Slope | -0.065 | A/B=1.03 | [0.71,1.29] | 0.338 |
| | | Paired t-test | High speed at 100% contrast | 0.111±0.008 | Low speed at 100% contrast | 0.122±0.006 | | | 0.230 |
| Fig.S1eii | Large pupil vs. Small pupil RT (n=39 mice) | Permutation | Large pupil Slope | -0.066 | Small pupil Slope | -0.060 | A/B=1.09 | [0.74,1.39] | 0.290 |
| | | Paired t-test | Large pupil 100% contrast | 0.114±0.007 | Small pupil 100% contrast | 0.133±0.008 | | | 0.023 |
| **Imaging** | | | | | | | | | |
| Fig.S4b | Linear/threshold vs. hyperbolic ratio fit of contrast response function (n=204 cells) | Paired t-test | Hyperbolic RMSE | 0.043±0.002 | Linear/Threshold RMSE | 0.043±0.002 | | | 0.498 |
| Fig.S4b | Linear/threshold vs. hyperbolic ratio fit of contrast response function (n=204 cells) | Paired t-test | Hyperbolic Adjusted $R^2$ | 0.54±0.02 | Linear/Threshold Adjusted $R^2$ | 0.67±0.02 | | | <0.0001 |
| Fig.3b | CPn vs. CSt visually-responsive neurons (n=6 mice, 391 cells vs. 6 mice, 332 cells) | Mann-Whitney U Test | CPn % Vis responsive | 34.1±10.6% | CSt % Vis responsive | 18.4±5.2% | | | 0.24 |
| Fig.3c | CPn vs CSt visual response slope (n=6 mice vs. 6 mice) | Mann-Whitney U Test | CPn Slope | 0.26±0.09 | CSt Slope | 0.10±0.02 | | | 0.015 |
| Fig.3d | CPn vs. CSt coefficient of variation slope (n=6 mice vs. 6 mice) | Mann-Whitney U Test | CPn Slope | -3.40±0.42 | CSt Slope | -2.02±0.28 | | | 0.0411 |
| Fig.3e | CPn vs CSt noise correlation (9453 pairs vs. 2145 pairs) | Kolmogorov-Smirnov test | CPn median | 0.126 | CSt median | 0.049 | | | <0.0001 |
| Fig.3f | CPn vs CSt locomotion modulation index on CRF Slope (138 cells vs. 66 cells) | One-sided t-test | CPn mean | 0.18±0.03 | Null | 0 | | | <0.0001 |
| | | One-sided t-test | CSt mean | 0.18±0.06 | Null | 0 | | | 0.0038 |
| | | t-test | CPn mean | 0.18±0.03 | CSt mean | 0.18±0.06 | | | 0.93 |
| Fig.4c | CPn correct vs. incorrect visual response slope (n= 6 mice) | Permutation | Slope correct | 0.26 | Slope incorrect | 0.18 | A/B=0.08 | [-0.08,0.07] | 0.0123 |
| Fig.4d | CSt correct vs. incorrect visual response slope (n=6 mice) | Permutation | Slope correct | 0.1081 | Slope incorrect | 0.1063 | A/B= 0.002 | [-0.10,0.05] | 0.4523 |
| Fig.4g (All) | CPn vs. CSt decoder d' (n=6 mice) | Mann-Whitney U Test | CPn decoder d' | 0.95±0.15 | CSt decoder d' | 0.30±0.20 | | | 0.009 |
| Fig.4g (High) Speed) | CPn vs. CSt decoder d' (n=6 mice) | Mann-Whitney U Test | CPn decoder d' | 1.00±0.16 | CSt decoder d' | 0.24±0.14 | | | 0.015 |
| Fig.4h | CPn single neuron decoder d' (n=138 cells) | One-sided t-test | CPn d' mean | 0.48±0.03 | CPn chance d' | -0.01±0.002 | | | <0.0001 |

| | | | | | | | | | |
|---|---|---|---|---|---|---|---|---|---|
| | CSt single neuron decoder d’ (n=66 cells) | One-sided t-test | CSt d’ mean | 0.07±0.04 | CSt chance d’ | -0.02±0.002 | | | 0.033 |
| | CPn vs. CSt single neuron decoder d’ (n=138 vs. 66 cells) | t-test | CPn d’ mean | 0.48±0.03 | CSt d’ mean | 0.07±0.04 | | | <0.0001 |
| | CPn vs. CSt single neuron decoder d' (n=6 mice) | Mann-Whitney U Test | CPn d’ animal-wise mean | 0.42+-0.08 | CSt d’ animal-wise mean | -0.10+-0.11 | | | 0.0087 |
| **Optogenetics** | | | | | | | | | |
| Fig.S5 a | CPn vs. CSt GFP-positive L5 cells (% NeuN-labeled cells) (n=6 vs 6 mice) | Mann-Whitney U Test | CPn proportion | 7.76±0.75% | CSt proportion | 7.28±0.42% | | | 0.589 |
| Fig.5c | CPn Laser on vs. Laser off Performance (n=11 mice) | Permutation | Laser on Rmax | 60.5% | Laser off Rmax | 80.2% | A/B=0.75 | [0.87,1.16] | <0.0001 |
| | | Paired t-test | Laser on 100% Contrast | 63.9±7.3% | Laser off 100% Contrast | 79.7±3.5% | | | 0.011 |
| Fig.5d | CSt Laser on vs. Laser off Performance (n=12 mice) | Permutation | Laser on Rmax | 79.3% | Laser off Rmax | 77.7% | A/B=1.02 | [0.91,1.09] | 0.325 |
| | | Paired t-test | Laser on 100% Contrast | 85.2±4.6% | Laser off 100% Contrast | 80.7±4.6% | | | 0.256 |
| Fig.S5 b | CPn Laser on vs. Laser off UR Amplitude (n=11 mice) | Paired t-test | Laser on UR | 0.91±0.02 | Laser off UR | 0.89±0.02 | | | 0.178 |
| | CSt Laser on vs. Laser off UR Amplitude (n=12 mice) | Paired t-test | Laser on UR | 0.91±0.01 | Laser off UR | 0.92±0.01 | | | 0.748 |
